# Characterization of human embryonic stem cells in animal component-free medium

**DOI:** 10.1101/2020.12.14.420984

**Authors:** Masakazu Machida, Rie Abutani, Hiroshi Miyajima, Tetsuji Sasaki, Yoshiko Abe, Hidenori Akutsu, Akihiro Umezawa

**Affiliations:** Center for Regenerative Medicine, National Center for Child Health and Development Research Institute, Tokyo, 157-8535, Japan; Cell Culture Business Promoting Division, Kyokuto Pharmaceutical Industrial Co., Ltd., Ibaraki, 318-0004, Japan; Business Strategy Division, SEKISUI MEDICAL CO., LTD., Tokyo, 103-0027, Japan

**Author notes:** **Corresponding author** Tetsuji Sasaki, Cell Culture Business Promoting Division, Kyokuto Pharmaceutical Industrial Co., Ltd., 3333-26, Asayama, Kamitezuna, Takahagi-shi, Ibaraki, 318-0004, JAPAN.

**Keywords:** Embryonic stem cells, Animal component-free, Feeder free, Single cell culture

## Abstract

Clinical use of human embryonic stem cells (ESCs) as a raw material requires good manufacturing practice-compliant axillary materials such as culture medium. To this end, animal components should not be used and contamination of virus/bacteria/fungus and allergens are a concern. In addition, animal components such as albumin and fetal bovine serum pose difficulties such as a lot-to-lot variation. However, only a limited number of animal component-free media have been developed to date. In this study, we investigated whether SEES2 ESCs can be stably propagated for 16 passages (54 population doublings) over a period of 60 days in a newly established Stem-Partner^®^ ACF medium. SEES2 ESC maintained their intact karyotype, i.e. 46,XX, and their undifferentiated phenotypes after long-term culture. An in vitro differentiation assay revealed that SEES2 ESCs exhibited multipotency, i.e. endodermal, mesodermal and ectodermal differentiation. Subcutaneous implantation of SEES2 ESCs generated mature teratomas without malignant transformation. These results show that SEES2 ESCs in the Stem-Partner^®^ ACF medium can be used to establish master cell banks for future regenerative medicine as well as other ESCs in the previously reported culture medium.

## Introduction

hESCs have the potential to differentiate into several types of cells and tissues in living organisms, and their capacity has attracted great interest in the field of regenerative medicine [1,2]. In general, pluripotent stem cells, including hESCs, are maintained in a heterologous or homologous protein-containing medium using mouse fetal fibroblasts (MEFs) as a feeder layer, which are derived from heterologous animals [3-5]. However, the clinical usage of hESCs for regenerative medicine requires high standards of safety and quality. Animal-derived components may pose a risk of viral infection or allergen-induced anaphylaxis to patients. Furthermore, animal-derived materials tend to show lot-to-lot variation in composition and activity, which may make reproducibility difficult. Therefore, it is necessary to eliminate animal-derived components from culture conditions to increase the safety and quality of regenerative medicine products.

The SEES2 cell line, an hESC lines, has been established at the National Center for Child Health and Development (NCCHD) for clinical applications and regenerative medicine research [6]. The SEES cell lines are presently being applied in clinical practice, including successful clinical trials in congenital urea cycle disorders [7,8]. It is necessary to establish a safer and more stable method of culturing SEES2 cells for clinical applications in the future. Akutsu and his research group evaluated a xenogeneic-free culture system using SEES2 cell line and successfully developed a safe culture system for hESCs by using gamma-irradiated knockout serum replacement and pharmaceutical-grade recombinant basic fibroblast growth factor (bFGF) in the medium [9]. They also reported that a feeder-free culture system using commercially available xenogeneic-free culture medium was also able to provide stable culture. In this study, we aimed to establish a complete animal component-free culture system by using Stem-Partner^®^ ACF medium, a medium free of animal-derived components, and vitronectin, a scaffolding material.

## Results

### Stable culture of SEES2 cells using the Stem-Partner^®^ ACF medium

We used SEES2 cells which were maintained on human mesenchymal stem cells (hMSCs) as a feeder layer with medium containing animal-derived components. We cultured SEES2 cells in the Stem-Partner^®^ ACF medium (animal component-free) under feeder-free and single cell culture conditions for 60 days. The cells showed stable growth during culture through 16 passages (Figure 1A). The morphology of the cells remained intact and did not show any significant changes even at later passages (Figure 1B). Immunocytochemistry confirmed that SEES2 cells were positive for pluripotency markers such as OCT3/4, Nanog, SOX2, and TRA1-60 (Figure 1C). Q-banding showed that SEES2 cells had intact karyotype, i.e. 46,XX (Figure 1D). These results indicate that SEES2 cells retain pluripotency under animal components-free and feeder-free conditions without chromosomal abnormality.

**Figure 1.**
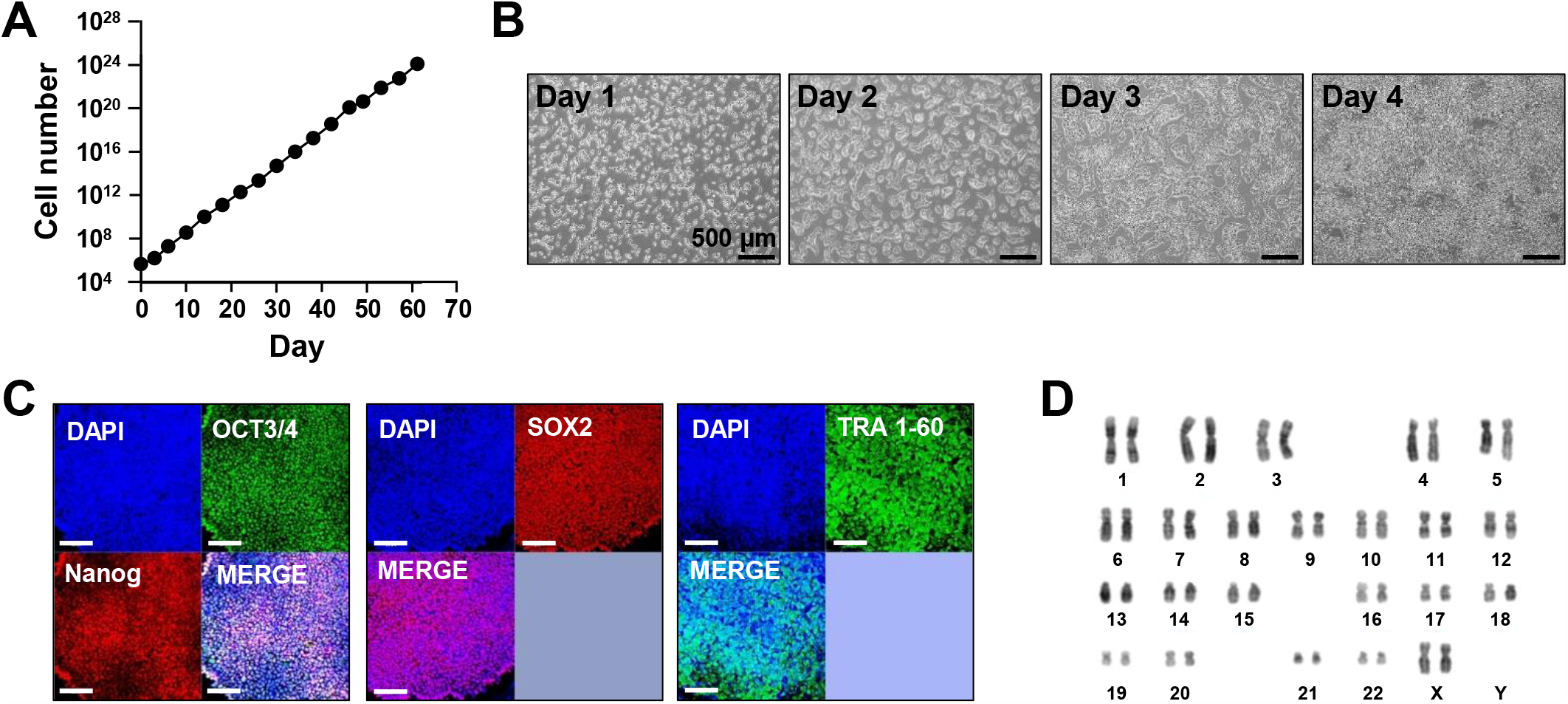
Characterization of SEES2 cells maintained under animal component-free conditions. A) Growth curve of SEES2 cells. SEES2 cells proliferated stably for 16 passages after the medium was changed to Stem-Partner^®^ ACF medium. B) Microphotographs of SEES2 cells from day 1 to day 4 of culture after 16 passages with Stem-Partner^®^ ACF medium. The SEES2 cells on each day were morphologically intact and did not show any differentiation. Scale bars are 500 μm. C) Immunocytochemical analysis of SEES2 cells after 16 passages with Stem-Partner^®^ ACF medium. Expression of pluripotency markers: OCT3/4, Nanog, SOX2, and TRA1-60, were observed. Scale bars are 100 μm. D) Karyotyping of SEES2 cells after 16 passages with Stem-Partner^®^ ACF medium. Karyotyping of SEES2 cells showed normal karyotype: 46,XX.

### Evaluation of pluripotency by embryoid body assays *in vitro*

To evaluate whether SEES2 cells maintained their pluripotency *in vitro*, we performed embryoid body (EB) assays. EBs differentiated from SEES2 cells expressed markers specific for three germ layers: tubulin beta III (TUJ1) (ectoderm), α-smooth muscle actin (SMA) (mesoderm), and α-fetoprotein (AFP) (endoderm) (Figure 2A). These results indicate that SEES2 cells cultured under animal component-free and feeder-free conditions maintained pluripotency.

**Figure 2.**
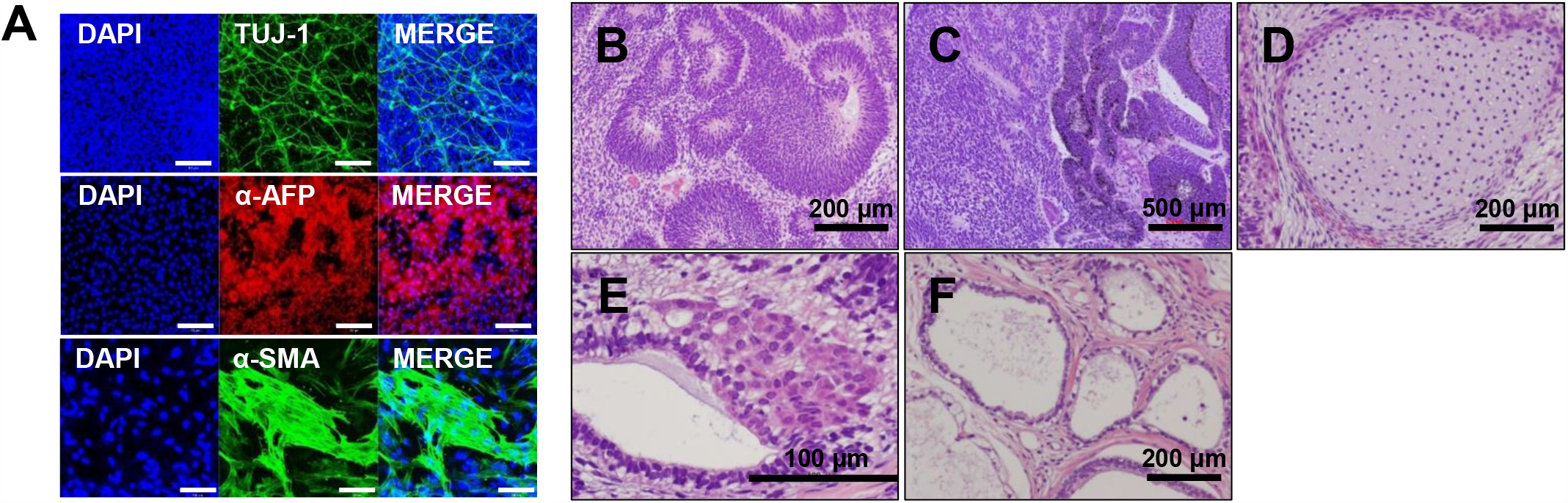
Differentiation analysis of SEES2 cells maintained under animal component-free conditions *in vitro* and *in vivo*. A) Immunohistochemical analyses of embryoid bodies differentiated from SEES2 cells after 16 passages with Stem-Partner^®^ ACF medium. Expression of germ layer-specific markers: TUJ1 (ectoderm), αSMA (mesoderm), and AFP (endoderm) were observed. Scale bars are 100 μm. B-F) Hematoxylin and eosin staining of teratomas generated by SEES2 cells *in vivo*. Immature teratomas differentiated from three germ layers were observed: B) neural tube-like structures consisting of immature neural tissue (ectoderm), C) melanocytes (ectoderm), D) cartilage (mesoderm), E) glandular structures consisting of a columnar epithelium (endoderm), F) glandular structures consisting of a cuboidal epithelium (endoderm).

### Evaluation of pluripotency by teratoma formation assay *in vivo*

To confirm whether SEES2 cells had the capacity to differentiate into specific tissues, we performed teratoma formation assays by implantation of SEES2 cells in the subcutaneous tissue of immunodeficient BALB/c AJcl-*nu/nu*mice. We observed the formation of ectoderm-derived tissues: neural tube-like structures consisting of immature neural tissue and melanocytes (Figure 2B, C), mesoderm-derived tissue: cartilage (Figure 2D), and endoderm-derived tissues: glandular structures consisting of a columnar epithelium or a cuboidal epithelium (Figure 2E, F). These teratomas did not show obvious malignant features, such as fetal carcinoma, choriocarcinoma and squamous cell carcinoma. These results indicate that SEES2 cells cultured under animal component-free and feeder-free conditions maintain pluripotency because benign teratomas formed by the differentiation of SEES2 cells contained derivatives shown characteristics of three germ layers.

## Discussion

As research into regenerative medicine using human embryonic stem cells and its application in clinical practice continues to advance, there is a growing need for regenerative medical products with a high level of safety and quality. One of the important indicators for assessing the safety and quality of regenerative medical product is whether or not animal-derived components are used. Another issue that arises when considering the clinical application of regenerative medical products is how to prepare large quantities of cells quickly. SEES2 cells, a strain of human embryonic stem cells, are starting to be used in clinical practice [7,8]. Akutsu et al. have studied a culture system without animal-derived components, and have reported that it is possible to culture cells stably in that culture system [9]. In the present study, we used Stem-Partner^®^ ACF medium, a commercially available culture medium that does not contain any animal-derived components, to investigate whether stable long-term culture of SEES2 cells is possible.

We confirmed that newly established culture system using Stem-Partner^®^ ACF medium and Vitronectin could easily condition SEES2 cells maintained on hMSCs as a feeder layer with medium containing animal-derived components. The SEES2 cells maintained in this culture system had normal chromosomes and remained undifferentiated after 16 passages. It was also shown that these SEES2 cells had pluripotency by EB assay and teratoma formation assay. These results indicate that we succeed in stable long-term culture. We additionally compared the growth rate of SEES2 cells in this culture system with a commercially available feeder-free culture system. As a result, no significant difference was observed in the growth rate in the logarithmic growth phase (data not shown). This result indicates that this culture system can provide as many cells as commercially available feeder-free culture systems.

We therefore consider that regenerative medical products manufactured using Stem-Partner^®^ ACF medium, which does not contain animal-derived components, can increase safety and quality when used in human clinical applications. Moreover, the Stem-Partner^®^ ACF medium can be expected to provide a high degree of reproducibility and stability in research applications because there is little lot-to-lot variation caused by animal-derived components. In addition, Stem-Partner^®^ ACF medium is compatible with the performance of metabolic analysis in that it does not contain proteins such as albumin or serum. The first challenge for the future will be the application of Stem-Partner^®^ ACF medium to other ESCs and three-dimensional cultures. Currently, there are many types of pluripotent stem cell lines in use. By accumulating experience in culturing other pluripotent stem cell lines and patient-derived induced pluripotent stem (iPS) cells using Stem-Partner^®^ ACF medium, we will be able to realize safe medical treatment and highly reproducible research. We would like to contribute to the development of the field of regenerative medicine by demonstrating the usefulness of the Stem-Partner^®^ ACF medium-based culture system for mass production using automated cell culture platforms and for 3D culture system that is close to physiological environments.

## Materials and methods

### Ethics

The ethics committee of the NCCHD specifically approved this study. The culture of hESC lines were performed in full compliance with “the Guidelines for Utilization of Human Embryonic Stem Cells (Notification of MEXT, June 7, 2017; Notification of MHLW, November 28, 2017; Notification of MEXT and MHLW, October 7, 2019)”.

### Animal component-free, feeder-free culture of SEES2 cells

SEES2 ESCs on γ-irradiated mouse embryonic fibroblasts were used for experiments using Stem-Partner^®^ ACF medium. Six-well plates (Thermo Fisher Scientific; 140675) were used as culture vessels. The 6-well plates were previously coated with Vitronectin-N (Thermo Fisher Scientific; A14700). Y-27632 (Fujifilm Wako, Tokyo, Japan; 259-00613) at a final concentration of 10 μmol/L and recombinant human full-length bFGF (#PHG0261) at a final concentration of 100 ng/mL were added to Stem-Partner^®^ ACF medium (Kyokuto Pharmaceutical Industrial Co., Ltd. Tokyo, Japan; #551-28143-9) just before use. SEES2 cells were suspended in prepared Stem-Partner^®^ ACF medium and diluted at 2.0 × 10^5^ or 5.0 × 10^5^ cells/2 mL. Diluted SEES2 cells were dispensed into Vitronectin-N-coated plates at 2 mL per well and cultured at 37°C. At intervals of 24 h, the culture medium was replaced with Stem-Partner^®^ ACF medium without Y-27632. Cell passages involved suspension of cells using TrypLE Select (Thermo Fisher Scientific, #12563011) at intervals of three to four days, centrifugal washing, and subsequent re-seeding of cells using Stem-Partner^®^ ACF medium containing Y-27632.

Stem-Partner^®^ ACF medium contains 21 amino acids (L-alanine, L-arginine, L-asparagine, Laspartic acid, L-cysteine, L-cystine, L-glutamic acid, L-glutamine, glycine, L-histidine, Lisoleucine, L-leucine, L-lysine, L-methionine, L-phenylalanine, L-proline, L-serine, L-threonine, L-tryptophan, L-tyrosine, and L-valine) as well as vitamins (ascorbic acid derivative, cobalamin, biotin, folic acid, I-inositol, niacinamide, d-calcium pantothenate, pyridoxine hydrochloride, riboflavin, thiamine hydrochloride, α-tocopherol and 4-aminobenzoic acid). It also includes trace elements, fatty acids and recombinant human growth factors. All ingredients used in Stem-Partner^®^ ACF medium are of non-animal origin. The medium component is a synthetic product made from minerals, plants, or microbial fermentation products, and does not contain any animal-derived material. The recombinant growth factors were produced from genetically modified microorganisms and plant cells, and not from animal cells. The medium was manufactured under ISO 13485 in a GMP-compliant facility, and stable for 10 months at −20°C.

### Immunohistochemical analysis of stem cell and differentiated markers

Immunohistochemical analysis was performed as previously described [9].

### Karyotypic analysis

Chromosomal Q-band analysis was performed at the Chromosome Science Labo. Ltd., Sapporo, Japan.

### Differentiation assays in vitro and in vivo

Differentiation assays in vitro and in vivo were performed as previously described [9].

## Conflict of Interest

Hiroshi Miyajima and Tetsuji Sasaki are employees of Kyokuto Pharmaceutical Industrial Co., Ltd., and Yoshiko Abe is an employee of SEKISUI MEDICAL CO., LTD.

## Author contribution

A. U., H. A., and T. S. designed experiments. M. M., R. A., and H. M. performed experiments.

A. U., H. M., and T. S. analyzed data. H. M. contributed reagents, materials, and analysis tools.

H. M. and Y. A. discussed the data and manuscript. A.U. and T. S. wrote this manuscript.

## Acknowledgements

We would like to express our sincere thanks to C. Ketcham for English editing and proofreading, and to E. Suzuki for English writing and secretarial work.

